# Mitochondrial localization of SESN2

**DOI:** 10.1101/871442

**Authors:** Irina E. Kovaleva, Artem V. Tokarchuk, Andrei O. Zeltukhin, Grigoriy Safronov, Aleksandra G. Evstafieva, Alexandra A. Dalina, Konstantin G. Lyamzaev, Peter M. Chumakov, Andrei V. Budanov

## Abstract

SESN2 is a member of evolutionarily conserved sestrin protein family found in most of Metazoa species. SESN2 is transcriptionally activated by many stress factors including metabolic derangements, oxidants and DNA-damage. As a result, SESN2 controls ROS accumulation, metabolism and cell viability. The best known function of SESN2 is the regulation of mechanistic target of rapamycin complex 1 kinase (mTORC1) that plays the central role in the stimulation of cell growth and suppression of autophagy. SESN2 inhibits mTORC1 activity through interaction with the GATOR2 protein complex that suppresses an inhibitory effect of GATOR2 on the GATOR1 protein complex. GATOR1 inhibits mTORC1 through its GAP activity toward the small GTPase RagA/B which in complex with RagC/D proteins stimulate mTORC1 translocation to the lysosomes where this kinase is activated by small GTPase Rheb. Despite the well-established role of SESN2 in mTORC1 inhibition, the other SESN2 activities are not well characterised. We recently showed that SESN2 can control mitochondrial function and cell death via mTORC1-independent mechanisms and these activities might be explained by direct effects of SESN2 on mitochondria. In this work we examined mitochondrial localization of SESN2 and demonstrated that SESN2 is located on mitochondria and can be directly involved in the regulation of mitochondrial functions.

## Introduction

Sestrins belong to evolutionarily-conserved protein family found in most of species of animal kingdom [1]. While invertebrate genomes contain only one gene encoding sestrin, genomes of vertebrates contain three *SESN1-3* genes. Sestrins are stress-responsive genes that play a major role in the regulation of cell viability through control of reactive oxygen species and metabolism [1]. Although sestrins are not required for embryogenesis they support homeostasis suppressing accumulation of age-related damages in different tissues of organism. Thus, our studies demonstrate that inactivation of sestrins in drosophila leads to deterioration of muscle tissue and excessive accumulation of lipids and other metabolites [2]. Inactivation of sestrin (cSesn) in *C. elegans* shortens the lifespan of the animals and weakens animals’ resistance to stress [3, 4]. Moreover, inactivation of the members of sestrin family in mice cause insulin resistance, sensitize animals to infarct and support development some types of cancer [5–9].

SESN2 is the best-characterised sestrin family member. The *SESN2* gene is activated by several transcription factors such as the tumor suppressor protein p53, the regulator of antioxidant response NRF2, and the regulator of integrated stress response ATF4 [1, 10–12] that indicate the potential role of SESN2 in the regulation of cellular homeostasis under these stress conditions. Our earlier works demonstrated that SESN2 modulates cell viability and the outcome depends on the type of stress the cells are encountered [10, 11]. We demonstrated that sestrins protect from ischemia and oxidative stress but can support cell death in response to DNA-damage and pro-apoptotic cytokines [10–13]. One of the major functions of sestrins is suppression of mechanistic target of rapamycin complex 1 kinase (mTORC1) [14, 15]. Sestrins suppress mTORC1 via interaction with the GATOR2 protein complex, composed of proteins Mios, WDR24, WDR59, Seh1L and Sec13 [16, 17]. GATOR2 works through inhibition of the GATOR1 complex, containing proteins DEPDC5, NPRL2 and NPRL3, which works as a GTPase activating protein for small RagA/B GTPases [18]. RagA/B in complex with RagC/D proteins interact with mTORC1 and translocate it to the lysosomal surface where it is activated by the small GTPase Rheb [19]. The interaction between SESN1/2 and GATOR2 complex can be negatively regulated by amino acid leucine (Leu) that binds the Leu-binding domain of sestrins and disrupt the interaction between sestrins and GATOR2, preventing negative effect of sestrins on GATOR2 and inhibition of mTORC1 [20]. However, many types of stress may stimulate formation of SESN2-GATOR2 complexes by means of activation of the expression of sestrins and, possibly, via some posttranslational modifications [16, 21]. Although GATOR1 proteins play major role in the regulation of mTORC1, they are also involved in the regulation of mitochondrial homeostasis and cell death in response to DNA damage [22]. Autophagy play a major role in regulation of cell viability in response to stress and SESN2 support the activation of autophagy and one of the specific form of autophagy – mitophagy either through inhibition of mTORC1, or through some other mechanisms such as interaction with autophagy receptor SQSTM1/p62 and the E3-ubiquitin ligase Rbx1 [14, 23].

SESN2 may also be involved in the regulation of metabolism and cell death through control of mitochondrial functions. We have demonstrated previously that SESN2-deficient mouse embryonic fibroblasts have lower ATP production comparatively to wild type counterparts [12, 24]. Mitochondria is the major source of ATP in cells and SESN2-deficient MEF have lower respiration rate comparatively to control cells [12]. Although in our analysis by immunofluorescence ectopically-expressed Flag-SESN2 protein was most in cytoplasm [16], we propose that some of the activity of SESN2 such as regulation of mitochondrial respiration and cell death may be mediated by direct effects of sestrins mitochondria. In this work we performed analysis of intracellular localization of SESN2 in the normal and stress conditions and demonstrated that SESN2 can be co-localized with mitochondria potentially regulating the activity of this organelle.

## Materials and methods

### Cell culture conditions

Human lung adenocarcinoma A549 cells, human osteosarcoma U2OS cells or human lung fibroblasts immortalized by SV40 T antigen, MRC5/T, cells were maintained in High Glucose DMEM containing 25 mM glucose and 10% FBS (Hyclone) or in the galactose medium consisted of DMEM deprived of glucose supplemented with 10 mM galactose, 2mM glutamine (4 mM final) and 10% dialyzed FBS (Hyclone) until they reached 70-80% confluence. For glucose starvation cells were rinsed with glucose-free DMEM and further incubated in glucose-free DMEM supplemented with 10% dialyzed FBS for 12 h. The cells were kept in CO2 5% at 37°C.

### Preparation of mitochondrial (HM) and cytosolic/LM fractions

A549 or MRC5/T were harvested from 150 mm dishes and homogenized in 550 μl of Mitochondrial Isolation Buffer (MIB) (220 mM mannitol, 70 mM sucrose, 10 mM Hepes pH 7.5, 1 mM EDTA and 1% CompleteTM protease inhibitor cocktail (Roche)) by passing the cell mixture through a 26-gauge needle at 4°C (50-100 strokes). The homogenate was centrifuged at 1000 g for 10 min at 4°C to remove the nuclei and the unbroken cells. The centrifugation step was repeated and the supernatant was centrifuged at 9 000 g for 10 min at 4°C. The pellet was washed with 1ml of MIB for two times. Approx. 5 μg of protein of the post mitochondrial supernatant and the pellet (crude mitochondria) were loaded per lane of 10% SDS/PAGE gel for Western blot analysis.

### Western blot analysis

Approx. 5 μg of protein per lane was separated on 10% SDS/PAGE gel and transferred onto a Hybond ECL^®^ (enhanced chemiluminescence) membrane (Amersham Biosciences). After blocking with 5% milk for 1 h, the membrane was incubated overnight at 4°C with primary antibodies diluted 1:1000 followed by a further 1 h incubation in the corresponding horseradish peroxidase conjugated secondary antibody (Bio-Rad) at a 1:10000 dilution, and the signal was detected with the ECL^®^ Western blotting detection system (Amersham Biosciences). The following primary antibodies were used for immunoblotting: AKAP1 (D9C5), COX4 (3E11), Mios (D12C6), Sestrn-2 (D186) from CST (Cell Signalling Technology); SESN2 (41-K) from Santa Cruz; Anti-Actin (ab3280) from Abcam; GAPDH (Sigma-Aldrich, G9545).

### Immunoprecipitation

Mitochondria isolated from 3 confluent 150 mm dishes were resuspended in the 200 μl of lysis NP40 buffer (0.3% NP40, 150mMNaCl, 50mMTris pH 7.5) supplemented with protease and phosphatase inhibitors (cOmplete Roche), incubated for 20 min at +4°C and centrifuged at +4°C 10,000 RPM for 15 min. SESN2 antibodies (Santa Cruz Biotechnology 41-K) or AKAP1 antibodies and 10 μl of protein A/agarose (Pierce) 50% slurry in the lysis buffer were added to the supernatant and then the mixture was incubated for 4 h on a rotating platform. The beads were washed 4 times with 1 ml NP40 buffer and reconstituted in the 50 μl of 2X electrophoresis loading buffer, and then the probes were heated at 1000C for 5 min, centrifuged 1 min at maximum speed and loaded on a gel for subsequent Western blot analysis.

### Proteinase K treatment

Isolated mitochondria were resuspended in MIB and treated with Proteinase K (50μg/ml) for 20 min on ice. The reaction was stopped by incubation with PMSF (1mM) for 10 min on ice and the probes were analyzed by Western Blotting.

### Fluorescent microscopy

To determine mitochondrial co-localization of SESN2, U2OS cells stably expressing GFP tagged SESN2 were exposed to 10 μM FCCP treatment for 24h and before the fluorescent microscopy procedure were incubated with 100 nM MitoTracker Red (CMXRos, Invitrogen) in DMEM for 30 min at 37°C to stain mitochondria.

### Analysis of respiration in *C. elegans*

#### Nematode strains

The wild-type WT and the cSesn^-/-^ (RB2325, genotype ok3157) animals with a 535 bp deletion in exon 3 of cSesn gene were purchased in the Caenorhabditis Genetics Center (CGC) of University of Minnesota and maintained on the sterile NGM (Nematode Groth Medium) containing 2% agarose, 0.3% NaCl, 1 mM MgSO_4_, 1 mM CaCl_2_, 0.25% 1 M phosphate buffer, 0.3% bactopeptone, and 0.05% cholesterol as described [4]. A night culture of *E. coli* (strain OP50, Uracil auxotroph) was spread out on agar beds of Petri dishes and the bacteria were grown overnight at 37°C. All experiments with nematodes were carried out at 20°C.

#### Synchronization of nematodes

To prepare a synchronous culture of nematodes, adult animals were treated with a solution containing 1% NaClO and 0.5 M KOH for 10 minutes. The released nematode eggs were washed twice in M9 medium [4] and incubated overnight at a temperature of 20°C in M9 medium. The next day, nematodes at the L1 stage were placed onto Petri dishes with seeded OP50.

#### Sample preparation

Synchronized L4 animals were rinsed from OP50 plates with M9 medium, washed twice, and rested in M9 medium for 20 minutes to remove bacteria from their guts to avoid the potential effects of bacteria on oxygen consumption rate (OCR) in *C. elegans*. Further, nematodes were resuspended in unbuffered EPA water (60 mg MgSO4×7H2O, 60mg CaSO4×2H2O, 4 mg KCl, ddH2O to 1L) one worm per microliter of the solution (confirmed by counting the number of worms in 20 μl drops). 75 microliters of the solution containing worms were then pipetted into each well of a 24-well Seahorse utility plate using tips rinsed in 0.1% Triton X-100 to prevent worms from sticking. The final volume of each well was then brought to 525 μl with unbuffered EPA water. Four wells in an assay were left as blanks. 75μL unbuffered EPA water with 80 μM FCCP (3.2 % DMSO), and 320 mM sodium azide were then loaded into the injection ports of the Seahorse cartridge. During Seahorse assay, after injection, each drug solution was diluted by eight times and final concentrations of drugs were 10 μM FCCP and 40 mM sodium azide.

### Analysis of mitochondrial respiration

The analysis of mitochondrial respiration in worms was done as described previously [25]. Seahorse software WAVE were set up for 16 cycles, consisted of a one mix cycle for oxygenation of the wells (1 min), a wait period to make worms settled down (3 min), and measurement of oxygen levels (3min). For initial determination of the basal OCR eight cycles were performed followed by injection with FCCP or azide. After the injection 8 more OCR measurements cycles were conducted. All OCR values were normalized per worm number as counted under microscope. Treatment worms with the mitochondrial uncoupler FCCP and complete respiratory inhibitor sodium azide was used for the determination of maximal respiratory capacity, spare respiratory capacity, and non OXPHOS OCR.

### Statistical analysis

Data was processed using STATISTICA software package (http://www.statistica.io). The comparisons of the two mean values was estimated using one-tailed Student’s t-test, *p≤0.05, **p≤0.01, ***p≤)0.001.

## Results

### Purification of SESN2 in mitochondrial fraction

To analyze localization of SESN2 we performed differential centrifugation of cellular lysates to get mitochondrial (heavy membrane fraction) and cytosolic/light membrane fraction as previously described [26]. As stress conditions may affect localization of the protein, we analyzed lysates from the cells kept in control condition, incubated in glucose-free medium or incubated in the medium where glucose was replaced for galactose. Both stress conditions lead to deficiency in energy production, however glucose starvation can also cause ER stress and oxidative stress [1]. Western blot analysis of cytosol fraction and membrane fraction demonstrated that considerable portion of the SESN2 protein was co-purified with the heavy membrane fraction enriched in mitochondria (Fig 1A). We also observed that the amount of SESN2 protein co-purified with membrane fraction was increased in stress conditions that may indicate that SESN2 is accumulated on the mitochondrial membrane to regulate the activity of the organelles or cell death. The analysis of samples indicates that the quality of our preparation was quite high as we were not able to observe the mitochondrial protein COX4 in the cytosolic fraction and the cytosolic protein GAPDH in the mitochondrial fraction (Fig 1A). As we used one tenth of the mitochondrial fraction and one hundredth of the cytosolic fraction for the loading on the gel, we can conclude that approximately equal amount of protein can be found in the cytoplasm and on the mitochondria according to the immunoblotting data (Fig 1A).

**Figure 1.**
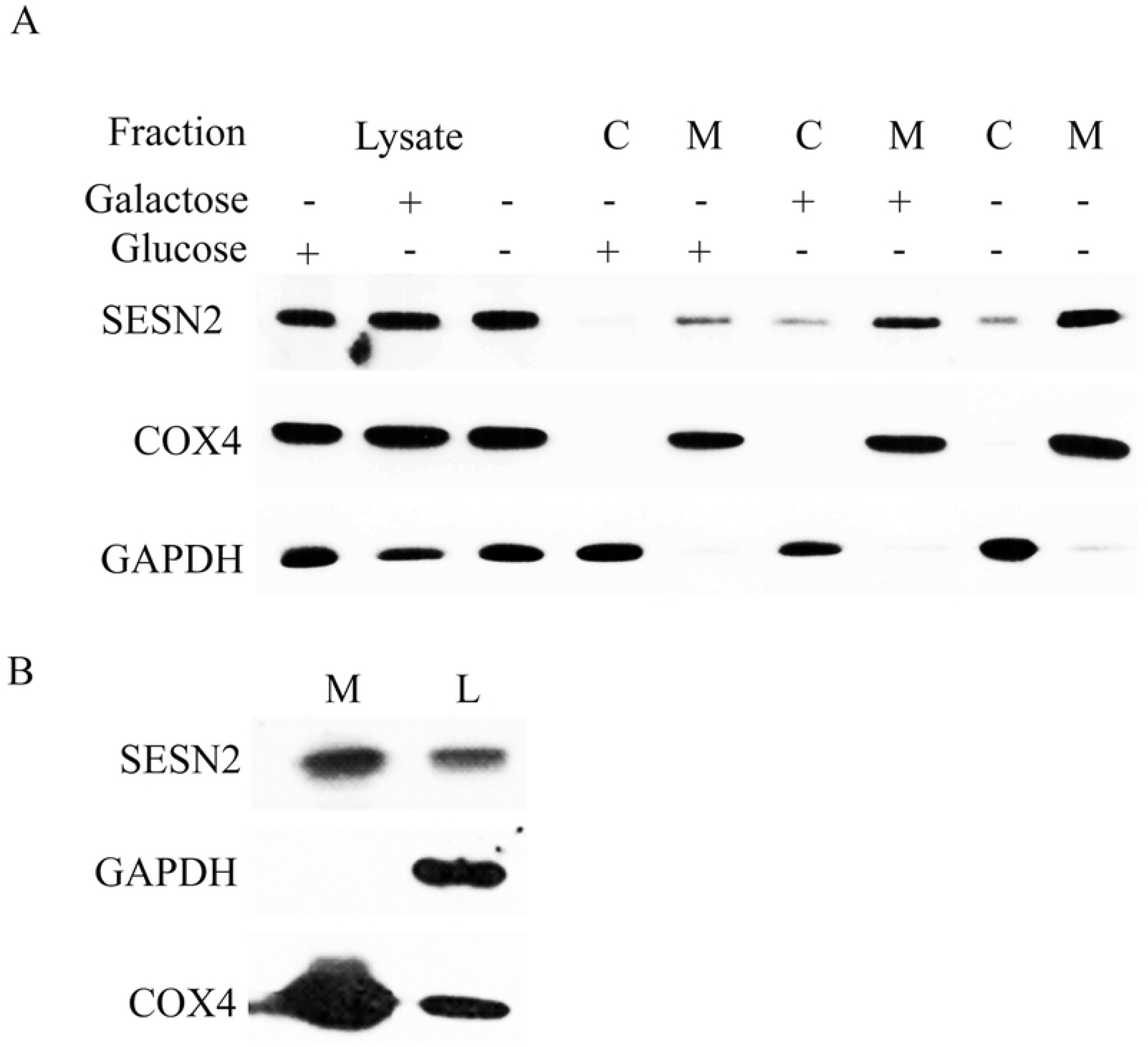
SESN2 is co-purified with mitochondria. (A) A substantial amount of endogenous SESN2 is detected in the fraction of crude mitochondria. A549 cells grown in indicated conditions were fractionated by differential centrifugation to get crude mitochondria fraction (heavy membrane, HM) and cytosolic/light membrane (LM) fraction. Equal protein amounts (5 μkg) of the whole cell lysate, cytosolic/LM fraction (C), and HM fraction (M) were analyzed by immunoblotting with anti-SESN2 antibodies to detect endogenous SESN2. The mitochondria inner membrane protein COX4 and cytosolic protein GAPDH were used as positive and negative controls of mitochondrial and cytoplasmic samples, respectively, to define mitochondria localized proteins. A549 cells were incubated in glucose-rich media, in the media where glucose was replaced for galactose or incubated in glucose-free media. (B) SESN2 is detected in mitochondria purified with magnetic beads bearing an antibody against mitochondrial outer membrane protein TOMM22. Mitochondria were isolated from A549 cells grown in galactose-containing media. M – purified mitochondria, L – whole cell lysate. 5 μg of the fraction protein was analyzed by immunoblotting with anti-SESN2, anti-GAPDH, and anti-COX4 antibodies.

Although differential centrifugation allows us to obtain enriched mitochondrial fraction, it can also contain some contaminants from other membrane organelles. To perform more accurate purification of mitochondrial fraction to confirm mitochondrial localization of SESN2, we performed high affinity purification of mitochondria using magnetic beads bearing antibodies against mitochondrial outer membrane protein TOMM22. The effectiveness of this approach was verified in several recent studies and is considered to be of full value an alternative to ultracentrifuge method [27, 28]. We observed that considerable amount of SESN2 was co-purified with the mitochondrial fraction supporting the previous observation that the protein is associated with mitochondrial fraction (Fig 1B). The high quality of purification of mitochondrial fraction is supported by the fact that we observe strong enrichment of mitochondrial protein COX4, but do not detect cytosolic GAPDH protein in the mitochondrial fraction (Fig 1B).

### SESN2-GFP fused protein is co-localized with a mitochondrial marker

To confirm that SESN2 can be located on mitochondria, we generated lentiviral expression construct where SESN2 is fused with fluorescent GFP protein as previously described [11]. U2OS cells were infected with the SESN2-GFP expressing recombinant lentivirus and treated with mitochondrial uncoupler FCCP to disrupt mitochondrial potential and inhibits mitochondrial respiration. To analyze localization of the SESN2-GFP protein in normal conditions or upon treatment with FCCP, the cells were incubated in the presence of Mitotracker Red and localization of the protein was assessed by fluorescent microscopy. A considerable portion of SESN2-GFP was co-localized with the mitochondrial marker regardless of FCCP treatment indicating that considerable amount of SESN2-GFP protein is located on mitochondria and this effect does not depend on the mitochondrial respiration (Fig 2).

**Figure 2.**
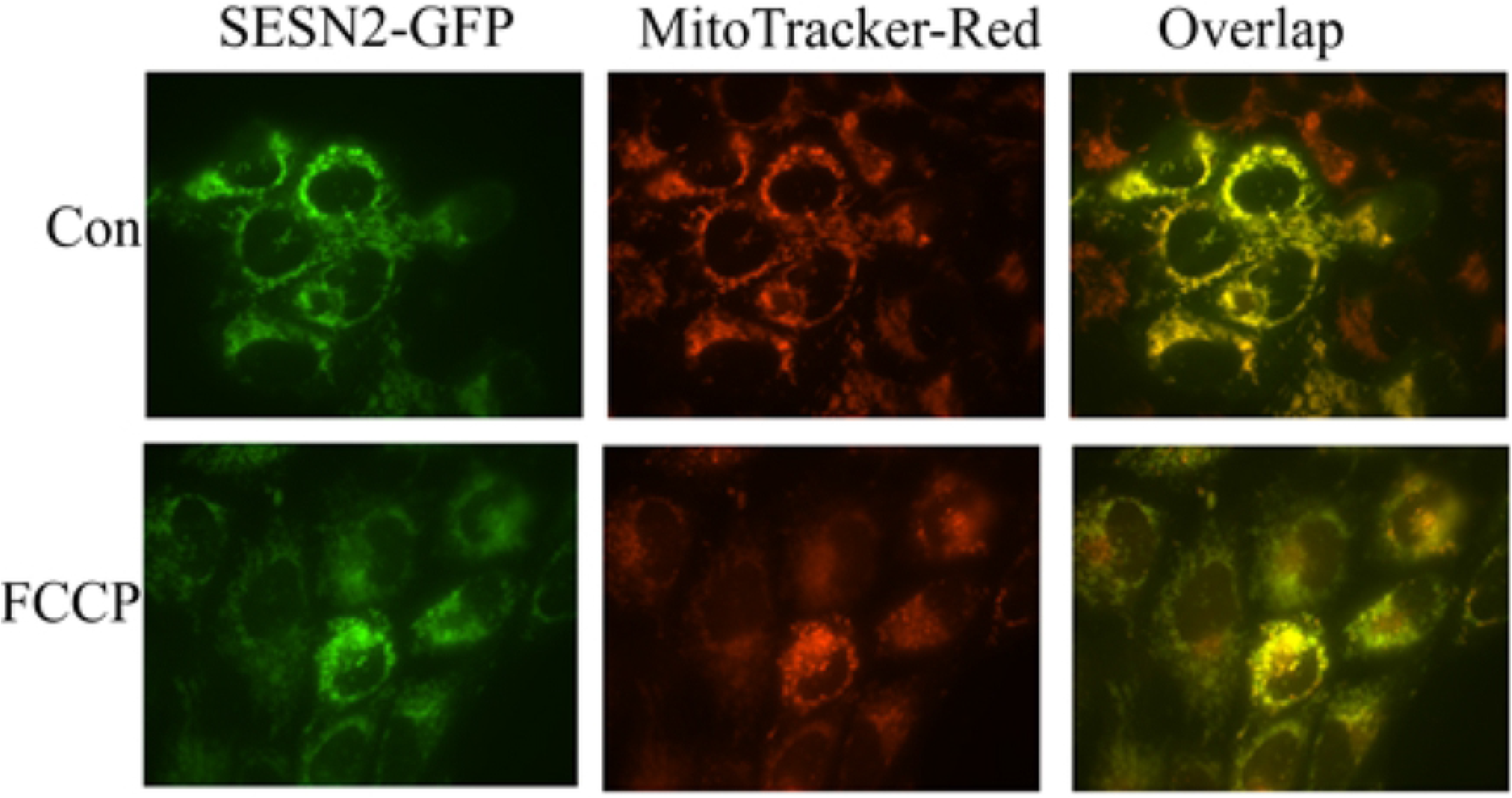
SESN2-GFP colocalizes with mitochondria. Confocal analysis of U2OS cells stably expressing fusion SESN2-GFP protein under basal conditions and upon treatment with 10 μM FCCP for 24 h. Mitochondria were stained with MitoTracker Red.

### SESN2 is co-localized with GATOR2 complex on mitochondria

We and others have previously shown that GATOR2 is the key partner of sestrins, however the precise localization of this protein complex in the cell is not well-established [16, 17]. We analyzed whether Mios, a component of the GATOR2 complex, can be detected in the mitochondrial fraction. We observed that mitochondrial fraction contains Mios as well as SESN2 proteins (Fig 3A). We did not see any difference in the amount of Mios in the mitochondrial fraction from control and SESN2-deficient cells indicating that mitochondrial localization of Mios does not require SESN2 (Fig 3A).

**Figure 3.**
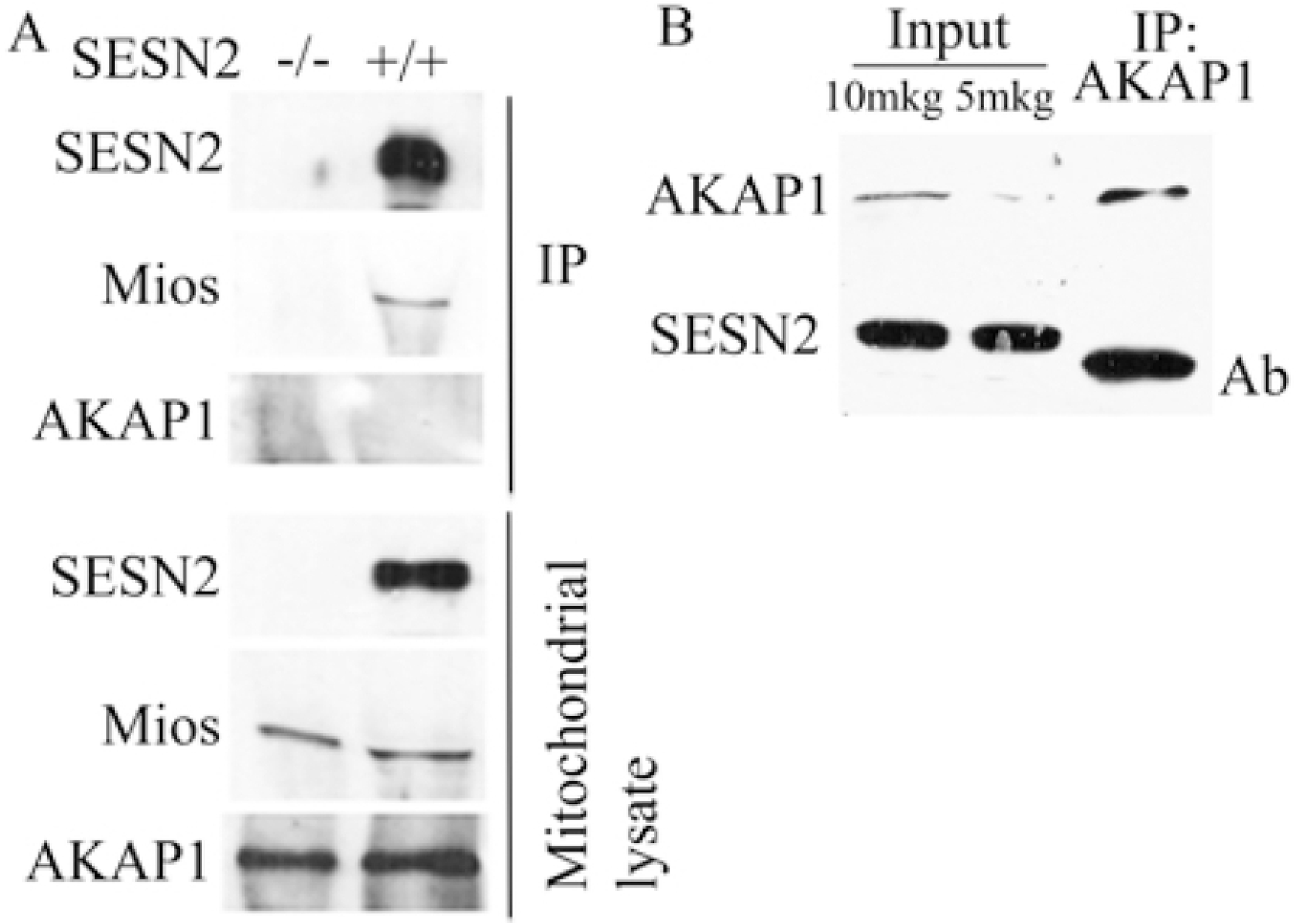
Mios coimmunoprecipitates with SESN2 from mitochondria lysates. (A) immunoprecipitation of S2 from mitochondria lysates. Mitochondria were isolated from control A549 cells or from A549 *SESN2^-/-^* cells after incubation in galactose media. SESN2-immunoprecipitates and mitochondria lysates were analyzed by Western blot with anti-SESN2, anti-Mios and anti-AKAP1 antibodies. (B) Immunoprecipitation of AKAP1 from mitochondrial lysates. Mitochondria were isolated from U2OS cells. Mitochondria lysate and AKAP1-immunoprecipitate were analyzed with anti-SESN2 and anti-AKAP1 antibodies.

To determine whether SESN2 interacts with Mios on the mitochondria, we immunoprecipitated endogenous SESN2 from mitochondrial fraction using anti-SESN2 antibody as previously described [16] and analyzed the presence of Mios in SESN2 immunoprecipitates. As a negative control we also analyzed mitochondrial lyzates from the cells where SESN2 was knocked out with CRISPR/Cas9 [29]. While we were able to detect Mios in the immunoprecipitates from control cells, indicating that these proteins interact on mitochondria, we did not observe any presence of Mios as well as SESN2 protein in the immunoprecipitated samples from SESN2-deficient cells (Fig 3A).

It was previously reported that SESN2 is capable to interact with mitochondria AKAP1 protein on mitochondria and control mTORC1 via AKAP1-dependent mechanism [30]. We analyzed whether AKAP1 is present in the mitochondrial fraction and in the SESN2 immunoprecipitates. While we observed notable amount of AKAP1 protein in mitochondrial fraction, we did not detect it in the SESN2 immunoprecipitates, indicating that it is unlikely that endogenous SESN2 interact with AKAP1 (Fig 3A). We also performed an analogous experiment using anti-AKAP1 antibodies but could not detect SESN2 in the AKAP1 immunoprecipitate (Fig 3B).

### GATOR2 affects mitochondrial localization of SESN2

We also determined whether mitochondrial localization of SESN2 depends on the GATOR2 complex. We generated cells where expression of either Mios or SehlL was silenced by shRNA lentivirus [16]. Both shRNA strongly downregulate expression of Mios (Fig 4) as it was previously reported [31]. In the cells where expression of GATOR2 proteins was silenced by shRNA we observed that amounts of SESN2 in mitochondrial fraction were notably decreased indicating that GATOR2 proteins support mitochondrial localization of SESN2 (Fig 4).

**Figure 4.**
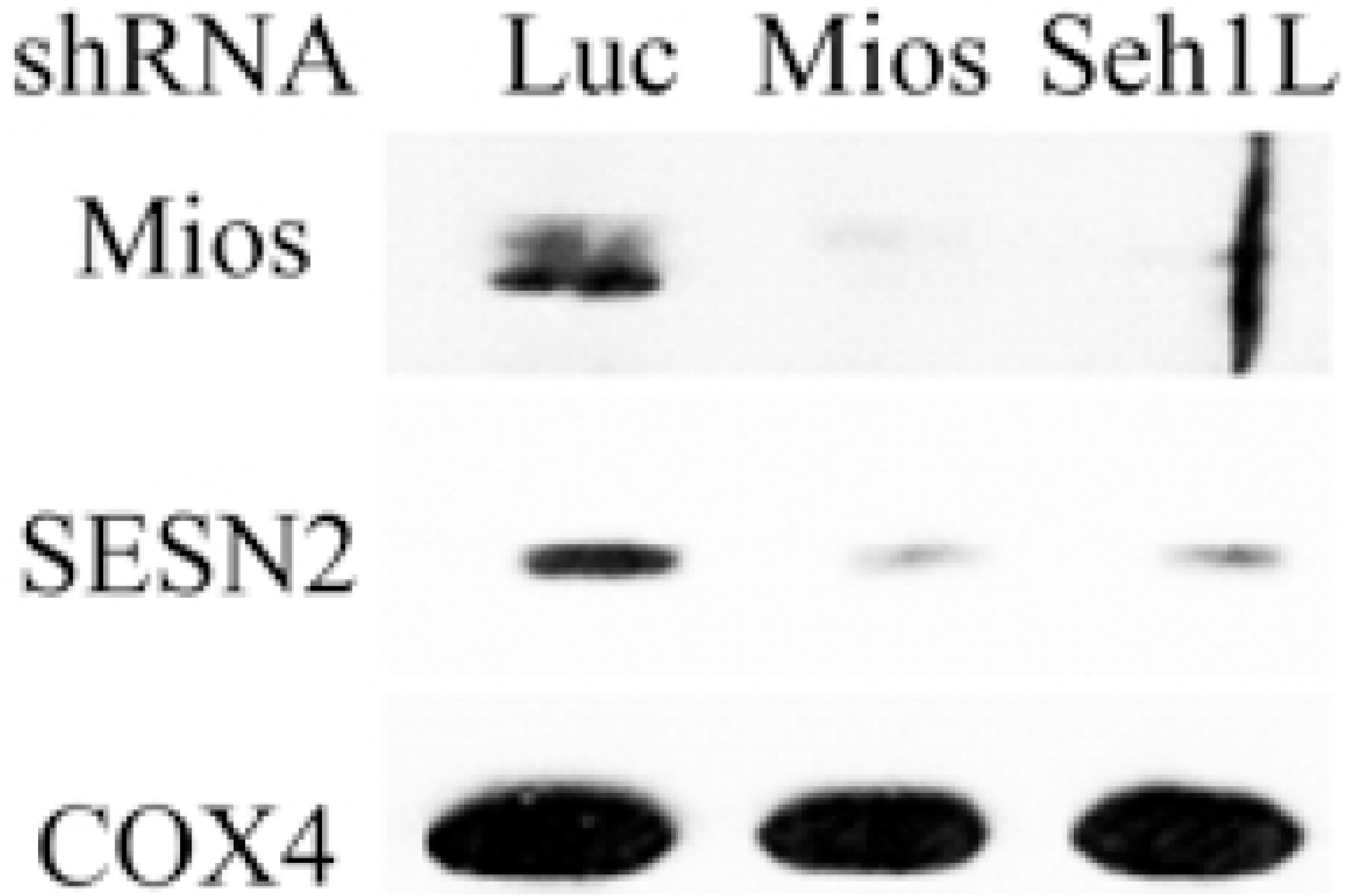
Downregulation of GATOR2 components Mios and Seh1L influence accumulation of SESN2 in the mitochondria fraction. Mitochondria were isolated from shLuc, shMios and shSeh1L A549 cells grown in galactose-containing media with magnetic beads conjugated to anti-TOMM22 antibodies.

### SESN2 is localized on the outer membrane of mitochondria or intermembrane space

To analyze whether SESN2 is transported inside of mitochondria or is located on the mitochondrial surface or intermembrane space we treated isolated mitochondria with proteinase K. We analyzed the presence of SESN2 in the untreated and proteinase K treated mitochondria and observed that treatment with proteinase K led to the disappearance of SESN2 from mitochondrial fraction. Similar effect was observed for voltage-dependent anion channel 1 (VDAC1) protein located on the outer mitochondrial membrane [32]. At the same time, we did not observe any changes in the amount of cytochrome C oxidase subunit 4 (COX4), the component of respiratory chain, imbedded in the inner mitochondrial membrane [33]. Therefore, we concluded that SESN2 is located on the surface of mitochondria or in the intramembrane space (Fig 5).

**Figure 5.**
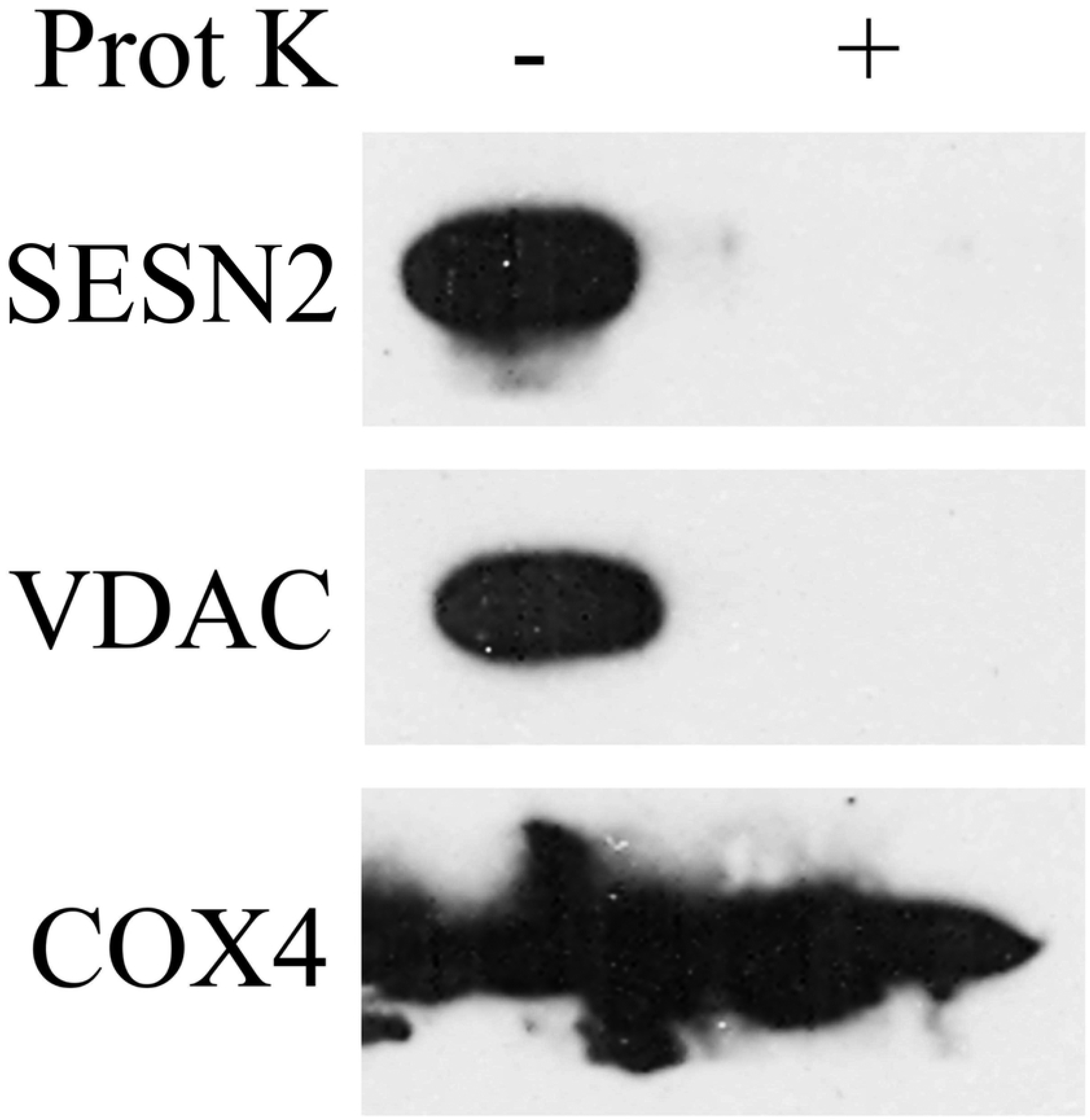
SESN2 is associated with outer mitochondrial membrane. Mitochondria fraction obtained from A549 cells was treated with 50 mg/ml proteinase K for 20 min on ice, and samples were subjected to Western blot analysis to detect endogenous SESN2. VDAC1 and COX4 were used as markers for mitochondrial outer membrane and inner membrane respectively.

### cSesn regulates mitochondrial respiration at *C. elegans*

Being associated with mitochondria, SESN2 and other members of sestrin family might be involved in the regulation of mitochondrial functions such as respiration, ROS production, and cell death [34]. To analyze the potential impact of sestrins on respiration, we analyzed an oxygen consumption rate in the cSesn-proficient and cSesn-deficient *C. elegans* that were generated in our recent studies [4]. We observed that the respiration levels and the levels of maximal respiration observed in the presence of an uncoupler FCCP were higher in cSesn-proficient animals as compared to WT controls (Fig 6A). Respiration in control and cSesn-null animals was squelched out by treatment with an inhibitor of mitochondrial respiration sodium azide [35], confirming the effects of cSesn on mitochondrial respiration.

**Figure 6.**
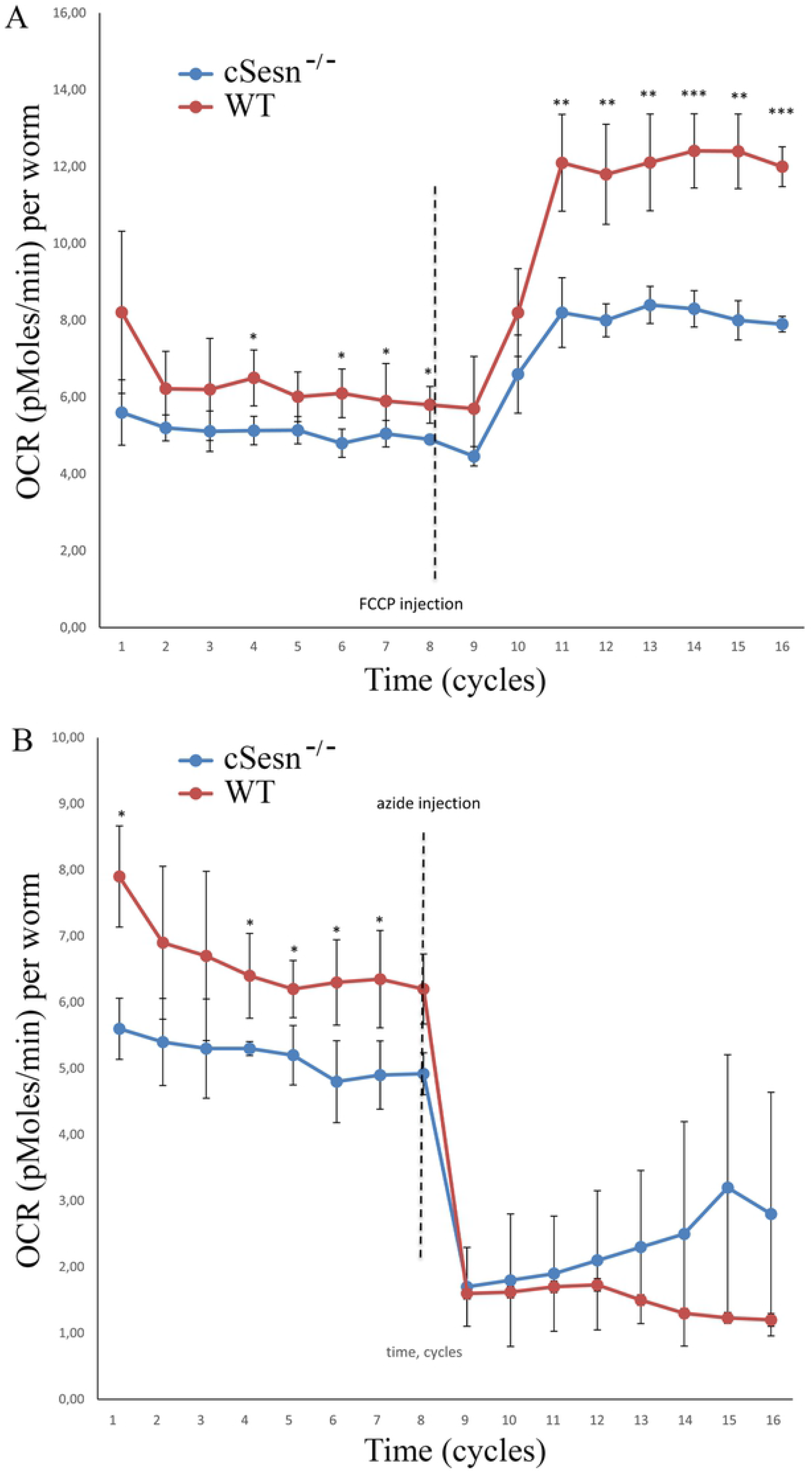
cSesn control mitochondrial respiration in *C. elegans*. (A) Analysis of basal and maximal oxygen consumption rate (OCR) in control (WT) and *cSesn^-/-^* animals. The OCR in the animals was analysed under control condition and after injection with FCCP. (B) Analysis of basal OCR and residual OCR after treatment with sodium azide. The comparisons of the two mean values was estimated using one-tailed Student’s t-test, *p≤0.05, **p≤0.01, ***p≤0.001.

## Discussion

Mitochondria play multiple roles in the cells being responsible for life-death decisions. Mitochondrial respiration is the major source of ATP but this process is also associated with leakage of electrons from the respiratory chain that lead to production of superoxide and other types of ROS [36]. Although at moderate levels ROS might play some signaling roles stimulating regulatory pathways involved in cell proliferation [37], exuberant accumulation of ROS causes damages of different types of macromolecules and compromise different physiological processes that can lead to senescence and cell death [38, 39]. Particularly, mitochondria is responsible for the induction of programmed cell death or apoptosis through accurately orchestrated mechanism involving release of mitochondrial proteins, formation of protein complexes called apoptosomes and activation of cysteine proteases – caspases, responsible for execution of apoptotic program [40]. Therefore, the timely and spatial control of mitochondrial function is extremely important for cells to ensure sufficient energy production, tune up metabolism, prevent ROS accumulation, and control cell death. One of the critical mechanisms of such regulation is mitophagy that monitor dysfunctional mitochondria and direct them to the degradation in the lysosomes [41]. There is a constant communication between mitochondrial and cytoplasmic proteins allowing cells to monitor dysfunctional mitochondria to be discarded via the autolysosomal pathway. The best-known process of monitoring of damaged mitochondria is driven by kinase PINK1 that is accumulated on the membranes of damaged mitochondria to recruit ubiquitine ligase Parkin involved in ubiquitylation of mitochondrial protein that label them for the consecutive degradation in the lysosomes. However, mitophagy is still observed in different tissues of PINK1-deficient animals indicating that this process can be also regulated through other, yet to be characterized mechanisms [42]. Some other proteins that normally function in different cellular compartments can also be associated with mitochondria and regulate mitochondrial function. For example this function was assigned to transcription factors STAT3 and p53 that are found associated with mitochondria where they regulate mitochondrial respiration or cell death, respectively [43, 44].

Stress-responsive SESN2 protein regulates variety of processes in cells including control of metabolism, ROS production and cell viability [1, 5]. Although it was well established that SESN2 and other members of the sestrin family are suppressors of mTORC1 that might work as leucine sensors, their role in cellular physiology is not limited by the regulation of mTORC1 and the mTORC1-dependent processes. We and others have shown that sestrins may modulate metabolism, cell death and autophagy/mitophagy in the mTORC1-independent manner [12, 13, 23, 45]. These data are supported by the observations that we often do not see difference in the mTORC1 activity between control and SESN1/2-deficient cells, while we still observe the clear impact of sestrins in the regulation of cell death and metabolism [12, 13]. It was previously suggested that sestrins undertake most of their functions in cytosol as ectopically-expressed Flag-labelled proteins were detected mostly in cytoplasm [16], however we were never able to specifically detect endogenous protein using available antibody when proper control SESN2-deficient cells were used. However, in a recent work it was demonstrated that SESN2 can be located in mitochondria by mean of interaction with AKAP1 protein and SESN2 can suppress the effects of AKAP1 on the mTORC1 activation [30]. AKAP1 is located on the outer mitochondrial membrane where it works as a scaffold protein that binds and regulates activities of several proteins involved in regulation of cellular signalling pathways such as protein kinase A, tyrosine kinase Src, and protein tyrosine phosphatase D1 [46]. As a result, AKAP1 is capable to control some mitochondrial functions such as respiration, Ca2+ homeostasis and cell death [46].

To resolve the inconsistency between our results and the observations by Rinaldi et al. [30] we performed several fractionation experiments to determine the localization of SESN2 in cells in the control conditions and upon incubation of cells with the glucose-free medium or with the medium where glucose in substituted by galactose. Both conditions affect ATP production causing cellular stress. Although cells survive on the medium supplemented with galactose, they intensify mitochondrial respiration to meet demand of ATP supply, while prolonged glucose starvation leads to necrotic cell death mediated by the stress of endoplasmic reticulum [12]. Using an approaches based on ultracentrifugation [26] we demonstrated that roughly a half of the SESN2 protein in the cell can be found in the mitochondrial fraction (Fig 1A). Interestingly, incubation of cells with glucose-free medium and galactose-containing medium increase the amount of protein associated with mitochondria. Our observations were confirmed by other studies where mitochondrial fraction was isolated by the mean of incubation of cellular lysates with beads coupled with the anti-TOMM22 antibody (Fig 1B). As TOMM22 is the protein located exclusively on the outer mitochondrial membrane, this approach allows us to isolate mitochondrial fraction of high purity [28]. The mitochondrial localization of SESN2 was also confirmed by our studies based on expression of SESN2-GFP fusion protein (Fig 2). The discrepancy with our previous data where the Flag-SESN2 protein was mostly observed in the cytoplasm can be explained by the application of a recombinant construct that makes expression of SESN2 extremely high permitting detection of the protein all over the cell and masking the portion of protein that can be co-localized specifically with mitochondria. In the current studies we used a recombinant construct that expresses protein under the control of relatively weak TP53 promoter that allowed us to keep the expression of SESN2 relatively low. Also the SESN2 protein is fused with GFP on C-terminus, while in the previous studies we used SESN2 linked with Flag epitope on the N-terminus that might also affect mitochondria translocation of SESN2 [16].

In our previous studies we purified the GATOR2 complex as major partner of sestrins [16] and here we determined that endogenous the SESN2 protein immunoprecipitated from mitochondrial fraction contained Mios, therefore, SESN2 might regulate mitochondrial functions via GATOR2 complex. In the same time, we were not able to detect AKAP1 in the complex with SESN2 or, alternatively, the SESN2 protein in the AKAP1-containing complex (Fig 3A-B). These data indicate that it is unlikely that AKAP1 is the bona-fide SESN2 partner on mitochondria and the previous data demonstrating the interaction between SESN2 and AKAP1 [30] can be explained by the overexpression of the SESN2 and AKAP1 proteins in HEK293 cells that can lead to non-specific binding between the proteins, especially taking into consideration that they might be located in close proximity on the outer mitochondrial membraine (Fig 5)[30]. Also AKAP1 was not detected in our mass spectrometry analysis of SESN2 protein complexes while we observed the presence of a considerable amounts of all GATOR2 proteins in the complex [16, 47]. However, we cannot rule out the possibility that SESN2 and AKAP1 can regulate each other through some indirect mechanisms and the relationshops between these two proteins are needed more elaborate characterization in the future. Also while it was reported that AKAP1 is capable to stimulate the activity of mTORC1 and SESN2 can counteract its effect, we were not able to observe the difference in the mTORC1 activity between control cells, cells with the deficiency of *SESN2, SESN1* or both *SESN1* and *SESN2* genes in cells incubated under normal conditions [29]indicating no impact of SESN1 and SESN2 in the control of mTORC1 despite the proposed role of AKAP1 in this process. We also demonstrated that while SESN2 does not affect GATOR2 localization on the mitochondria, GATOR2 facilitated mitochondrial localization of SESN2, supporting the idea that AKAP1 is probably dispensable for this process (Figs 3–4).

The function of GATOR2 on the mitochondrial membrane is not well-understood. Interestingly, the partner of GATOR2 - the protein complex GATOR1 was found associated with mitochondria where it might interact with the regulator of cell death, mitochondrial protein AIF [22]. Therefore, GATOR1 can support release AIF from mitochondria in response to some stress stimuli that may lead to nuclear translocation of AIF facilitating induction of cell death [22]. The SEACAT complex, homolog of GATOR1 in yeast, is also involved in the regulation of mitophagy in yeast and is involved in maintenance of mitochondria-vacuole contact sites [48]. These observations demonstrate that GATOR1 might have some mTORC1-independent functions on mitochondria, and GATOR2, being a bonafide GATOR1 interactor [18, 49] might be involved in the negative regulation of these functions. Therefore, through the control of GATOR2 SESN2 might regulate the GATOR1 functions on mitochondria. However, we cannot rule out the possibility that SESN2 might also regulate mitochondrial activities in a GATOR1/2-independent manner. We have previously shown that Sesn2 regulates mitochondrial respiration and ATP production in control conditions and in the conditions of glucose deprivation in mouse embryonic fibroblasts [12]. Interestingly, these effects were not dependent on the supply of intermediates mitochondria respiration and mitochondria size[12]. In these studies we extended our previous observation using cSesn-negative C. elegans as an alternative model where we demonstrated that inactivation of cSesn decrease mitochondrial respiration rate (Fig 6). The data obtained on the alternative model system may indicate that sestrins might play an evolutionarily-conserved role in the regulation of mitochondrial respiration. Although direct effect of sestrins on the regulation of the components of mitochondrial respiration chain can be considered, the alternative mechanism may involve mitochondrial quality control mediated by the regulation of mitophagy. Accordingly, the role of SESN2 in the support of mitophagy was recently reported [45, 50, 51]. These data are also supported by the activation of SESN2 in response to inhibitors of mitochondrial respiratory chain such as rotenone and piericidin A [16, 52] that might indicate that SESN2 might be required to provide support of mitochondrial function in the conditions of mitochondrial stress. Another potential mitochondrial function of sestrins might be the regulation of cell death. Accordingly, in our earlier studies we demonstrated that SESN2 regulated cell viability supporting cell death in response to DNA damage but suppressing cell death in response to ischemia and oxidative stress [10]. The detailed analysis of the role of sestrins on mitochondria in the future studies will allow us to better understand the role of sestrins in the control of cellular homeostasis. As mitochondrial disfunction underlies many human age-related diseases, some of the protective effects of sestrins against diabetes, infarct, cancer, and neurodegenerative diseases [5, 15] can be explained by their support of mitochondrial function and sestrins might be desirable target for development of antiaging drugs aimed to support mitochondrial activities.

## Authors contributions

Purification of mitochondrial fractions: IEK, AGE, and GS. Immunoprecipitation: IEK. Microscopy analysis: AVT and KGL. Construct preparation and lentiviral infection: AAD, IEK and PMC. Analysis of mitochondrial respiration: AOZ. Experiments’ design and paper preparation: AVB.

## Acknowledgements

The work was supported by the Grant 17-14-01420 from the Russian Science Foundation to AB, grant 075-15-2019-1660 from the Russian Federal Research Program for Genetic Technologies Development to PC. We also thank Nadushulya Pryadilova for help with the manuscript.

